# Investigating the risk of cardiac fibrosis due to heat-not-burn cigarettes through human cardiac stromal cells

**DOI:** 10.1101/2022.09.06.506632

**Authors:** Vittorio Picchio, Francesca Pagano, Roberto Carnevale, Alessandra D’Amico, Claudia Cozzolino, Erica Floris, Antonella Bordin, Leonardo Schirone, Wael Saade, Fabio Miraldi, Elena De Falco, Sebastiano Sciarretta, Mariangela Peruzzi, Giuseppe Biondi-Zoccai, Giacomo Frati, Isotta Chimenti

## Abstract

**Background:** The use of alternative smoking devices, such as heat-not-burn cigarettes (HNBC), is increasing on a global scale, and their impact on health is still uncertain.

**Objective:** To investigate the effects of circulating molecules in HNBC chronic smokers on the fibrotic specification and paracrine function of cardiac stromal cells (CSCs).

**Methods:** Resident CSCs were isolated from the atrial tissue of patients with cardiovascular diseases, and exposed to the serum of 60 young healthy subjects, stratified in exclusive HNBC smokers, traditional combustion cigarette (TCC) smokers, or non-smokers (NS) as reference.

**Results:** CSCs treated with TCC serum displayed impaired 3D growth and migration, as well as increased expression and/or release of pro-inflammatory and pro-fibrotic cytokines. Cells cultured with HNBC serum showed increased mRNA levels of pro-fibrotic genes, and reduced expression of the gap junction protein CX43. Nonetheless, both TCC and HNBC sera reduced the release of angiogenic and protective factors from CSCs. In fact, their paracrine support to tube-formation by endothelial cells and to preserved cell viability of cardiomyocytes in culture was significantly impaired. Treatment with the sera of both types of smokers also increased the expression of NOX isoforms and the release of H_2_O_2_ by CSCs.

**Conclusion:** The circulating molecules in the serum of chronic HNBC smokers induce fibrotic specification in CSCs. They also reduce the beneficial paracrine effects of stromal cells on endothelial cells and cardiomyocytes, albeit to a reduced extent for some features. These results point to a potential risk for atrial fibrosis development triggered by chronic HNBC use.

**CONDENSED ABSTRACT:** The use of alternative smoking devices, such as heat-not-burn cigarettes (HNBC), is increasing on a global scale, and their impact on health is still uncertain. We isolated human stromal cells from the atrial tissue of patients with cardiovascular diseases, and exposed them to the serum of young healthy subjects, that are exclusive HNBC smokers. Results showed significant alterations in the phenotype of CSCs exposed to HNBC serum, suggesting a specification towards fibrosis, reduced support to parenchymal cells, and increased oxidative stress production. Data point to a potential risk for atrial fibrosis development triggered by chronic HNBC use.

## INTRODUCTION

Traditional tobacco combustion cigarettes (TCCs) are main risk factors for lung cancer and cardiovascular diseases. Besides the role of TCCs in the etiology and progression of endothelial damage and atherosclerosis, many studies indicate an important causal role also in triggering myocardial fibrosis. This increases the risk of atrial fibrillation and promotes maladaptive cardiac remodeling mechanisms in chronic smokers (1,2). The main cell type responsible for fibrosis and remodeling in the heart are resident cardiac stromal cells (CSCs). These cells respond to pathological stimuli through activation from a primitive homeostatic state, and differentiate into collagen-producing cells, such as myofibroblasts (3,4). Other than direct fibrosis contribution by collagens deposition, CSCs also exert indirect effects through paracrine mechanisms on parenchymal and immune cells, as they influence cell survival and stress resistance, angiogenesis, and immune cell activation (3,5,6).

The use of alternative smoking devices, such as vaping electronic cigarettes and, more recently, heat-not burn cigarettes (HNBCs), is increasing dramatically on a global scale (7), but their effects on human health, particularly in the long term, are still uncertain (8). In fact, epidemiological studies on long-standing risks cannot be achieved yet, since these products are relatively new on the market. The recent SUR-VAPES-2 study (9) has showed how a single use of HNBC causes an acute adverse impact on circulating markers of oxidative stress, platelet function, flow-mediated dilation, and blood pressure. These effects, though, appeared reduced compared to a single use of TCC. Interestingly, the subsequent SUR-VAPES Chronic study (10) has reported that young healthy subjects (average age 30), having exclusively used HNBCs for at least 18 months, have increased oxidative stress, endothelial dysfunction, and platelet activation when compared with matched non-smokers. Most importantly, this study could not detect any differences in these parameters in comparison with chronic TCC smokers, suggesting a highly detrimental cardiovascular impact of HNBCs.

Parallel to clinical research on modified risk products, many studies have assessed their molecular and biological effects on different cell types, mainly considering endothelial and lung cells (11,12). Most of these studies, though, have been performed by direct exposure of cells to vaping e-cigarettes liquids or derived aerosols. Besides, the more recently introduced HNBCs are essentially unexplored. Moreover, these products could exert their effects in chronic smokers most likely through modification of the circulating molecules profile, particularly concerning interstitial cell types that are not directly exposed to smoke. Thus, the biological response of myocardial cell types to circulating signals in chronic HNBC smokers, and the overall molecular and cellular mechanisms of cardiovascular damage mediated by HNBCs are still unknown.

We hypothesized that the significant alteration of circulating signals caused by chronic use of HNBCs can induce a pro-fibrotic phenotypic shift in resident CSCs, reducing their homeostatic properties (e.g. migration, angiogenic and trophic support) while fueling detrimental paracrine and functional features (e.g. pro-fibrotic and pro-inflammatory). To this aim, we used resident primitive CSCs isolated from non-smoker patients with cardiovascular diseases, and exposed them to sera of patients enrolled in the SUR-VAPES Chronic clinical trial, including chronic HNBC smokers, chronic TCC smokers, and matched non-smokers.

## METHODS

Detailed methods and supporting data are available as online supplemental material.

### Human cardiac stromal cells culture

Human adult cardiac stromal cells (CSCs) were isolated from right atrial appendage tissue, as previously described (13,14), during clinically indicated procedures, after informed consent, under protocol 2154/15 approved by the Ethical Committee of “Umberto I” Hospital, “La Sapienza” University of Rome. In brief, tissue was fragmented, digested, and plated as explant cultures in dishes previously coated with Fibronectin (FN) (Corning) in complete explant media (CEM) formulated as follows: Iscove’s modified Dulbecco’s medium (IMDM) (Sigma-Aldrich) supplemented with 20% FBS (Sigma-Aldrich), 1% penicillin-streptomycin (Sigma-Aldrich), 1% L-glutamine (Lonza), and 0.1 mM 2-mercaptoethanol (Thermo-Fisher). Explant-derived cells were collected after 3 weeks by sequential washes with Ca2+-Mg2+ free PBS, 0.48 mM/L Versene (Thermo-Fisher) for 3 minutes, and with 0.05% trypsin-EDTA (Lonza, Basel, CH) for 5 minutes at room temperature. Collected cells were selected for a primitive non-activated phenotype by spheroid growth, as previously described (13,15), to deplete the culture from differentiated cells, such as myofibroblasts or smooth muscle cells; cells were then expanded as a monolayer on FN-coating in CEM. Three CSCs lines were used, each obtained from a non-smoker donor, free from dysmetabolic conditions (diabetes mellitus, or metabolic syndrome), and under beta-blockers therapy (13,15).

### Statistical analysis

For clinical record analyses, variables are reported as median (1st; 3^rd^ quartile) for continuous variables, and count/total (percentage) for categorical variables. Differences were computed with unpaired Mann-Whitney U test for continuous variables, and Fisher exact test for categorical variables using SPSS (IBM, version 25, USA). Significance was set at <0.05. For experimental data, results are presented as mean value±SEM, unless specified. For in vitro treatment results, inter-patient variability was controlled for by normalizing experimental results of each CSC line versus the NS control, and averaging fold-change modulations. Datasets were checked for significant outliers by Prism 8 software (GraphPad Software, San Diego, USA), which were excluded when present. Normality of data was assessed and significance of differences among groups was determined by Two-way ANOVA, with Bonferroni or Fisher LSD post-test, as appropriate, using Prism 8 (GraphPad Software). A p-value <0.05 was considered significant.

## RESULTS

### Retrospective analysis of CSCs isolated from chronic smokers

Since the features of primitive CSCs derived from chronic TCC smokers had never been described, as a preliminary step we retrospectively analyzed experimental data of resident CSCs isolated from a cohort of 38 patients undergoing cardiac surgery for cardiovascular diseases (15). A previous study from our group had evidenced a significant negative effect of chronic beta-blockers therapy (13) on the phenotype of resident CSCs isolated from this set of patients. Therefore, we only considered cells isolated from patients under beta-blocker therapy, and then stratified them based on smoking habits: non-smoker donors (never smoked, 20 CSC lines), and smoker donors (former or current, 18 CSC lines). The two cohorts were fairly homogeneous, except for a higher proportion of males, a slightly lower average age, and a higher proportion of coronary bypass surgery indication in the smokers’ group (Supplemental Tables 1 and 2). Available cell culture, flow cytometry, and gene expression data were evaluated. The analysis evidenced a significant impairment in a typical mesenchymal/stromal feature, that is spontaneous growth as 3D spheroids, in CSCs isolated from smoker patients versus non-smokers (Figure 1A). In detail, 15 out of 20 (75%) primary cell lines derived from non-smokers were able to spontaneously grow as spheroids, while only 6 out of 18 (33,3%) from smoker patients did. Moreover, a significant increase in the percentage of cells positive for the pro-fibrotic marker CD90 emerged in CSCs isolated from smokers compared to non-smokers (Figure 1B). Finally, gene expression analysis showed significantly higher levels of the myofibroblast markers *ACTA2*, and of the oxidative stress enzymes *NOX2* and *NOX5* in CSCs derived from smokers compared to non-smoker donors (Figure 1C).

**Figure 1.**
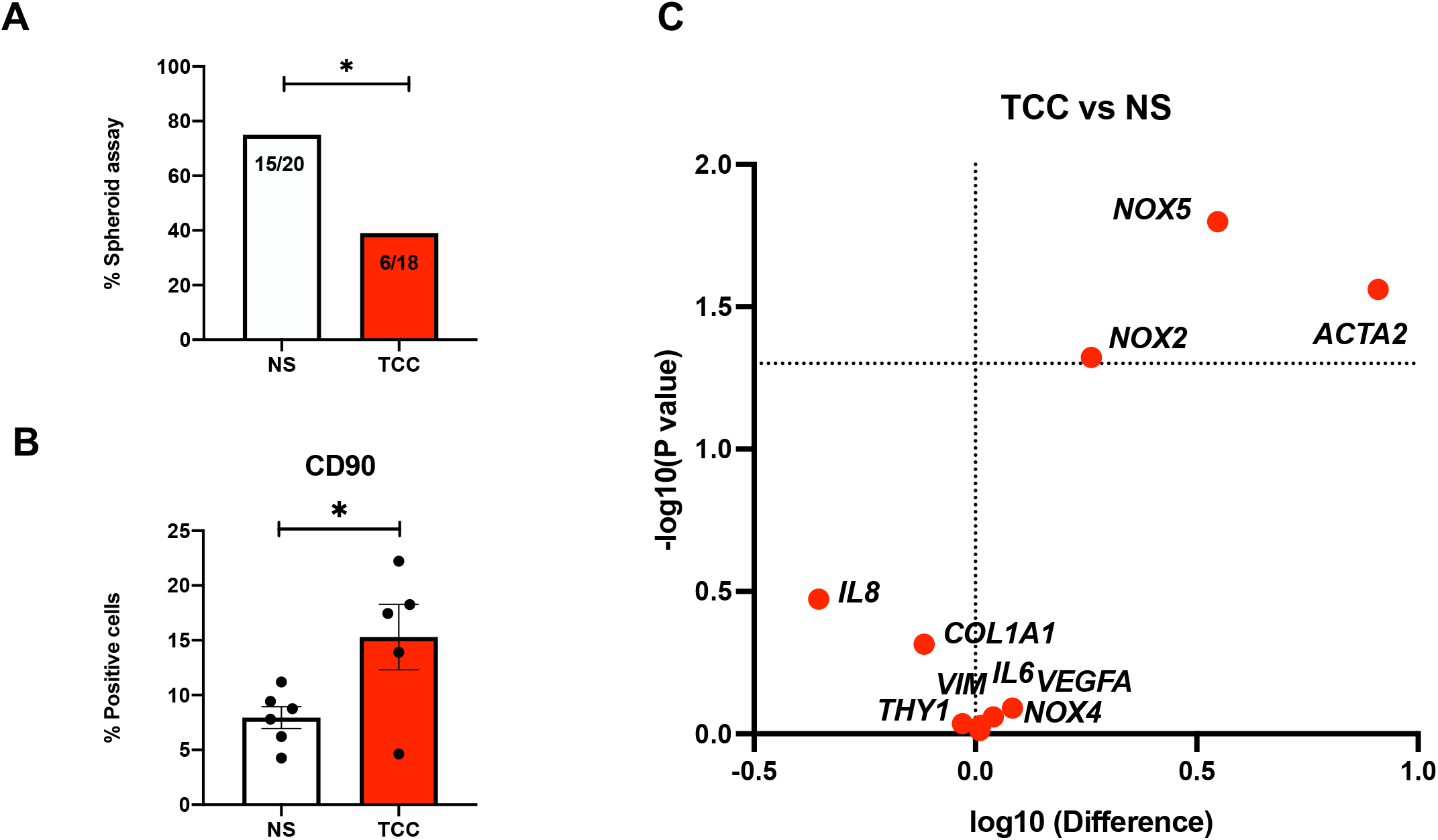
Retrospective analysis of the phenotype of CSCs isolated from smokers. **A)** The percentage of CSC lines able to successfully grow as spheroids, isolated from non-smokers (NS) versus tobacco combustion cigarette (TCC) smokers, is reported. Significance was computed by chi-squared test. **B)** CSCs isolated from smokers contained a higher proportion of CD90+ pro-fibrotic cells compared to those from non-smokers, as assessed by flow cytometry. N≥5. **C)** Gene expression levels (plotted as logarithmic difference in absolute expression of CSCs from smokers versus non-smokers) with significance are presented as volcano plot, showing how CSCs from smokers expressed significantly higher levels of the oxidative stress markers *NOX2* and *NOX5*, and of the myofibroblast markers *ACTA2*, compared to CSCs from non-smokers. N≥6. * = P<0.05.

### Cell phenotype after treatment with serum from TCC and HNBC smokers

We investigated the effects of *in vitro* exposure to the serum of chronic TCC or HNBC smokers on CSCs derived from atrial biopsies of 3 non-smoker patients undergoing elective cardiac surgery. We selected CSCs from three donors undergoing beta-blocker therapy (13), and free from diagnosis of dysmetabolic conditions (metabolic syndrome, or type 2 diabetes) (15). Clinical and surgical features of the 3 cell donors are reported in Table 1. Serum had been collected previously from young healthy non-smokers (NS), chronic heat-not burn cigarette (HNBC) smokers (mean exclusive use of 1.5±0.5 years), and chronic tobacco combustion cigarette (TCC) smokers (mean exclusive use of 7 years), enrolled in the SUR-VAPES Chronic study (10). Small identical volumes of serum from the 20 patients per group were pooled to create three serum lots used in all following experiments with CSC cultures. Each of the 3 CSC lines was then cultured up to 1 week in media supplemented with 10% serum lot of either NS, HNBC or TCC smokers.

**Table 1.** Clinical and surgical characteristics of the 3 selected biopsy donors for CSC isolation. They were selected among patients under beta-blockers therapy, non-smokers (never smoked), and free from diagnosis of dysmetabolic conditions (type 2 diabetes or metabolic syndrome).

|  | **L41** | **L45** | **L69** |
| --- | --- | --- | --- |
| **Age, years** | 68 | 73 | 78 |
| **Gender** | Male | Male | Female |
| **BMI, kg/m<sup>2</sup>** | 26.6 | 27.7 | 31.3 |
| **BSA, m<sup>2</sup>** | 1.9 | 1.9 | 1.9 |
| **Systolic BP, mmHg** | 130 | 120 | 120 |
| **Diastolic BP, mmHg** | 80 | 80 | 70 |
| **Heart rate, bpm** | 55 | 75 | 60 |
| **Atrial fibrillation** | No | No | Yes |
| **Previous MI** | No | Yes | No |
| **Recent MI (<30 days)** | No | No | No |
| **STS mortality risk score** | 0.93 | 1.69 | 2.18 |
| **Euroscore II** | 2.52 | 2.07 | 2.7 |
| **Type of surgery** | Coronary bypass | Coronary bypass | Heart valve surgery |
| **CPB time, min** | 121 | 107 | 180 |
| **Cross clamp time, min** | 96 | 87 | 140 |

BMI: body mass index. BSA: body surface area. BP: blood pressure. MI: myocardial infarction. STS: society of thoracic surgery. CBP: cardiopulmonary bypass.

The spontaneous capacity to grow in 3D was quantified, as a feature of mesenchymal phenotypes (16,17). The number of spheroids formed by plating the same cell number significantly decreased when CSCs were cultured in the TCC serum lot versus the NS lot (Figure 2A-B), with comparable spheroid size among groups (Figure 2C). This result suggests a reduction in a typical stromal feature in treated CSCs, consistent with the retrospective analysis (Figure 1A).

**Figure 2.**
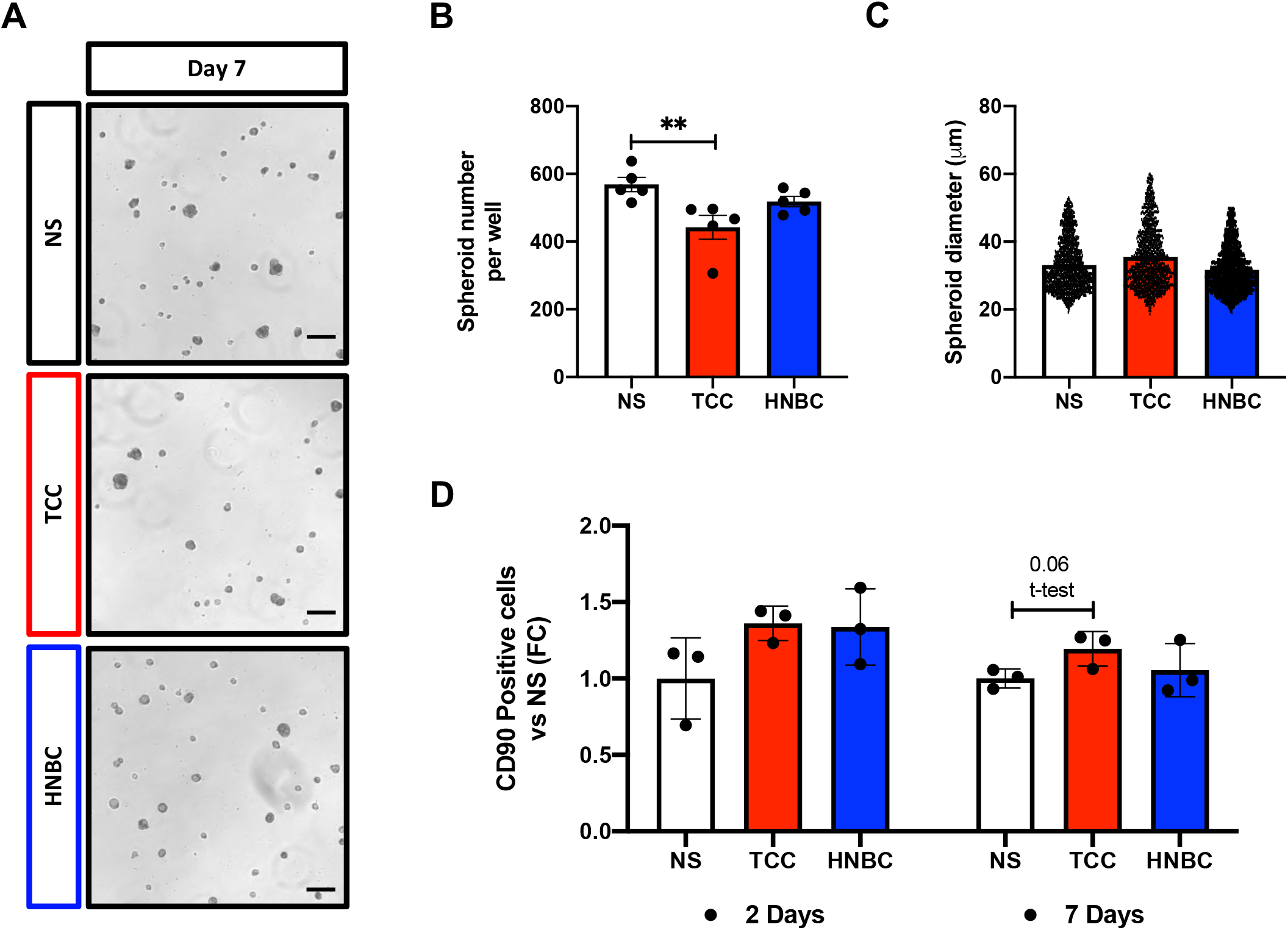
Spheroid growth and phenotype of CSCs after treatment with serum lots of smokers. **A)** Representative bright field microscopy images of spontaneous spheroid growth of CSCs treated with the serum lots of non-smokers (NS), heat-not burn cigarette (HNBC) smokers, or tobacco combustion cigarette (TCC) smokers. Scale bar = 100 μm. Spheroid number per well **(B)**, and average size of spheroids **(C)** were assessed for 3 CSC lines after 7 days of culture with the different serum lots. N (analyzed spheroids per well) >300. The percentage of CD90+ CSCs was quantified by flow cytometry after 2 and 7 days of culture **(D)**, and reported as fold change (FC) versus NS. N=3. * = P<0.05, ** = P<0.01, *** = P<0.001.

When cultured as monolayers, a trend towards increased percentages of cells positive for the fibrosis marker CD90 (p=0.06) was detectable by flow cytometry in CSCs cultured with the TCC serum lot compared to the NS lot (Figure 2D), consistent with the profile of CSCs directly isolated from chronic smokers observed from the initial retrospective analysis (Figure 1B). The gene expression profile of CSCs exposed for 48 hours to treatment with the 3 serum lots is presented in Figure 3. This analysis evidenced a significant up-regulation of the pro-inflammatory cytokines *IL6* and *IL8* in CSCs exposed to the TCC serum lot versus the NS lot (Figure 3A), as well as a significant up-regulation of the extra cellular matrix (ECM) remodeling marker *MMP1*, of the fibroblast activating cytokine *PDGFA*, and of the oxidative stress enzyme *NOX5*. We also detected a slight, but significant downregulation of type III collagen (*COL3A1*) and *NOX4* in cells treated with the TCC serum lot compared to the NS control (Figure 3A). On the other side, CSCs treated with the HNBC serum lot displayed significant upregulation of type I collagen (*COL1A1*), *THY1* (encoding for CD90), *PDGFA*, and of the oxidative stress enzyme *NOX2*, all versus treatment with the NS lot (Figure 3B). Conversely, *VEGFA, GJA1* (encoding for connexin 43), and *MMP1* were significantly downregulated after culture with the HNBC serum lot compared to the NS control (Figure 3B). When evaluating gene expression levels between TCC and HNBC serum lot treatments (Supplemental Figure 1), results showed significantly higher levels of *IL6, IL8, VEGFA* and *MMP1*, and significantly lower levels of *COL1A1* in CSCs cultured with the TCC serum lot versus the HNBC lot.

**Figure 3.**
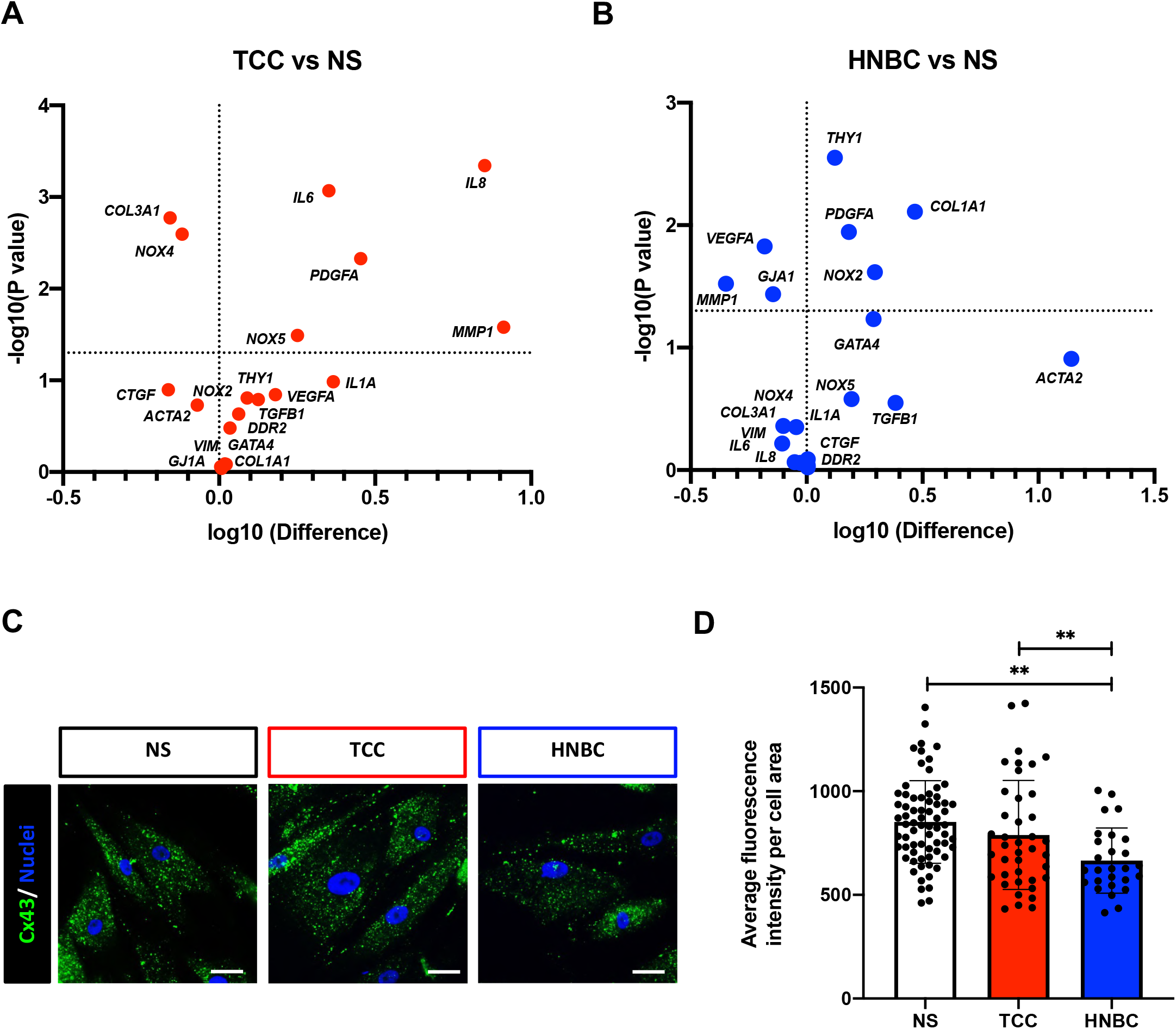
Gene expression analysis of CSCs after treatment with serum lots of smokers. The relative gene expression levels of several markers, with corresponding significance values, is plotted as volcano plot for the comparison of tobacco combustion cigarette (TCC)-versus non-smoker (NS)-serum lot treatments of CSCs **(A)**, and of heat-not burn cigarette (HNBC)-versus NS-serum lot treatments **(B)**. N=3. Representative immunofluorescence panels **(C)**, and quantification of average fluorescence intensity per cell area **(D)** of Connexin 43 (CX43) after 48 hours of culture of CSCs in the different serum lots. N (analyzed cells per condition) ≥28. ** = P<0.01. Scale bar = 50 μm.

Since connexin 43 is the main mediator of CSC electrical integration, and since its deranged modulation has been associated with atrial fibrillation (18,19), we further assessed its expression by immunofluorescence staining. Results showed a small, but significant reduction in the average fluorescence intensity per cell area in CSCs treated with the HNBC serum lot for 48 hours, compared to both NS and TCC lots (Figure 3C-D), consistently with the gene expression modulation.

Next, we analyzed the differential capacity of the three serum lots to stimulate migration in CSCs. We observed a significantly higher residual scratch area after 6 hours for cells treated with the TCC serum lot versus the NS lot (Figure 4 A-B). Cells cultured with the HNBC serum lot demonstrated a delayed migration compared to the NS group, albeit without reaching statistical significance. A higher residual area in CSCs exposed to the TCC serum lot was maintained up to 10 hours, instead the effect on migration of the HNBC serum gradually aligned with that of the NS control (Figure 4C-D). This suggests that serum from chronic TCC smokers has an impaired capacity of sustaining stromal cells migration, while that of HNBC smokers exerts an intermediate effect.

**Figure 4.**
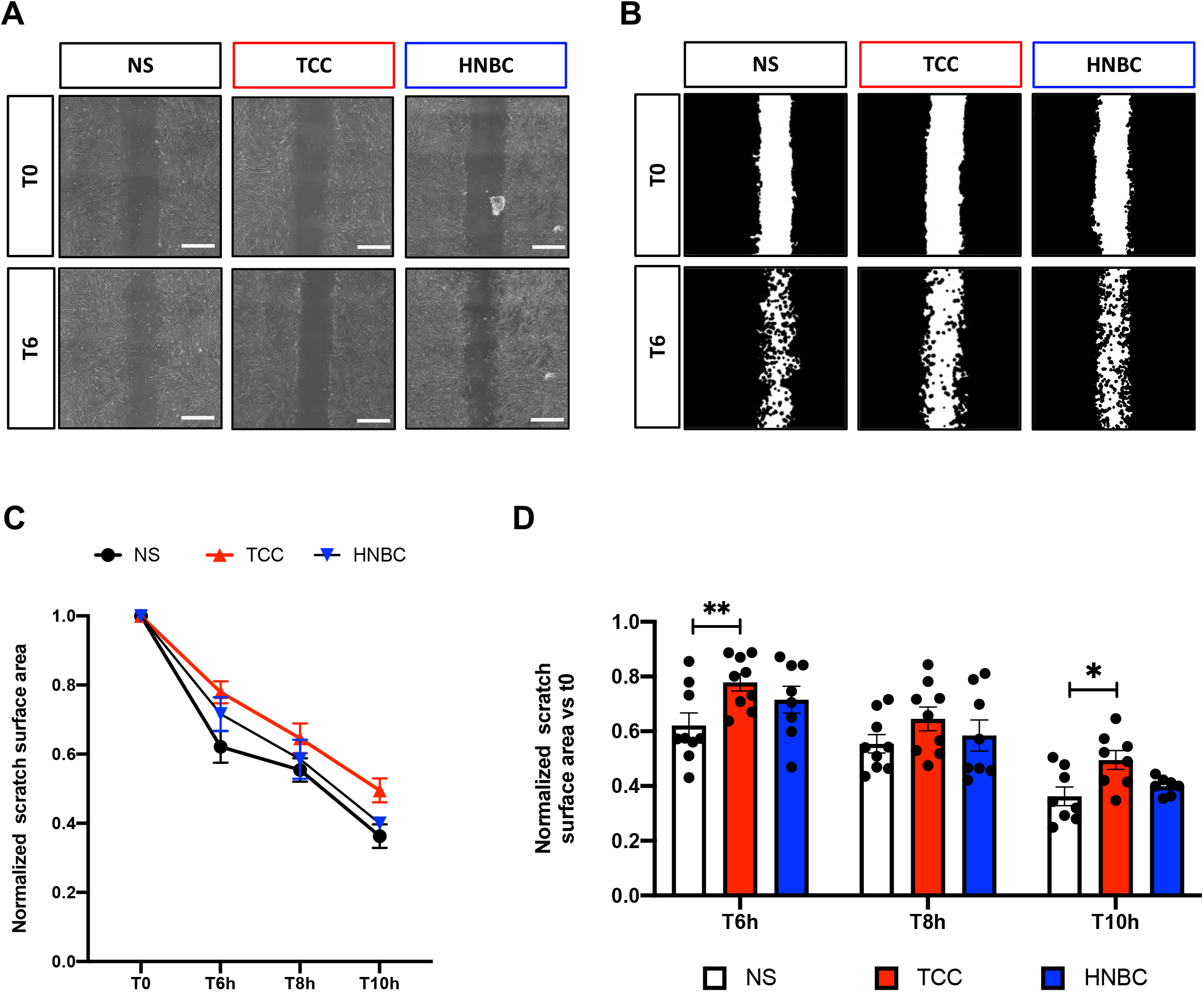
CSC migration capacity in response to serum lots of smokers. Representative bright field microscopy images **(A)** and relative binary masks obtained by the ImageJ macro **(B)** of the residual area of the scratch assay performed with the 3 CSC lines (each in experimental triplicate) cultured in serum lots from non-smokers (NS), tobacco combustion cigarette (TCC) and heat-not-burn cigarette (HNBC) smokers. The quantification of the residual surface area is plotted as normalized values in time **(C)**, as well as dot plots bar graph for each single time-point **(D)**. N=9. * P<0.05, ** P<0.01. Scale bar = 500 μm.

### Paracrine profile after treatment with serum from TCC and HNBC smokers

After 48 hours of CSC culture, human serum-supplemented media were removed, and replaced with 0.1% FBS-IMDM, which was then collected after 24 hours. These conditioned media (which contain only molecules released by CSC) were screened by protein arrays, and log2 optical density (OD) values for cytokines with OD≥0.2 were plotted as heatmap (Figure 5A). The analysis showed that several cytokines were similarly downregulated after exposure to serum lots from both types of smokers, including the anti-inflammatory and anti-fibrotic molecules adiponectin, CHI3L1 and GDF-15, and the angiogenic factor VEGF. Another set of cytokines, instead, including pro-inflammatory molecules such as VCAM, C5/C5A, PTX3, MIF, CFD, SDF-1, CD40L, MMP9, IL-17a, PAI1, CRP, Apoa1, and CXCL4, were released at higher levels only by CSCs previously cultured in the TCC serum lot, compared to the other groups. Interestingly, we detected an increased release of Vitamin D binding protein (VITDBP) and sex hormone binding globulin (SHBG) by CSCs cultured in both TCC and HNBC serum lots, compared to the NS control.

**Figure 5.**
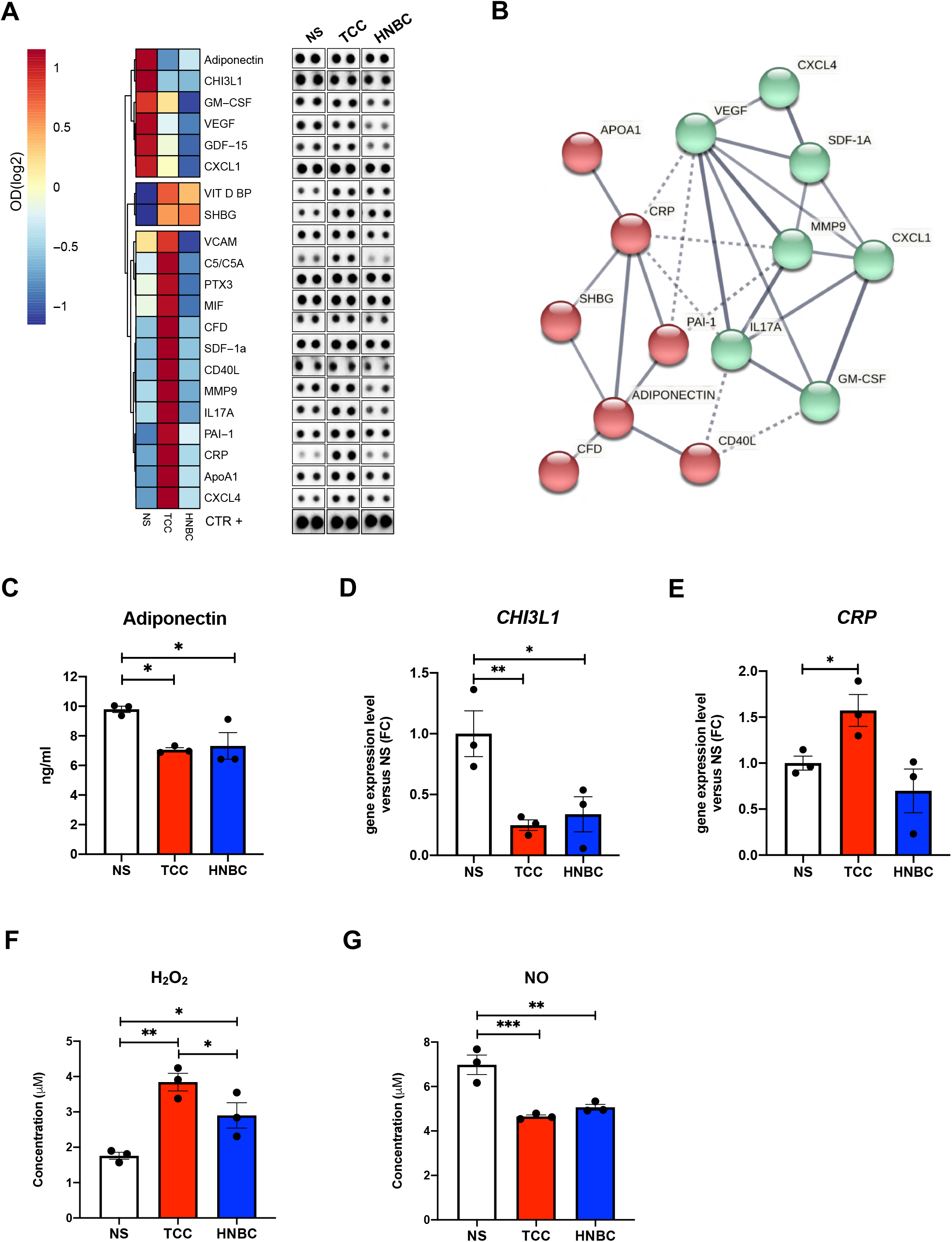
Profile of secreted molecules by CSCs after treatment with serum lots of smokers. Conditioned media (without human serum) was collected after 24 hours from CSCs previously treated for 48 hours with serum lots from non-smokers (NS), tobacco combustion cigarette (TCC) and heat-not-burn cigarette (HNBC) smokers, and screened by protein arrays; log2 optical density (OD) values were then plotted as heatmap, with corresponding representative blots shown beside **(A)**. Network of functional associations obtained from the STRING database **(B)**. Red e green colors identify the 2 clusters detected by k-means clustering. Dotted lines represent connections between cytokines in different clusters. Validation of the protein arrays was performed by ELISA for adiponectin **(C)**, and by RT-qPCR for *CHI3L1* **(D)** and *CRP* **(E)**. The concentrations of hydrogen peroxide (H_2_O_2_) **(F)** and nitric oxide (NO) **(G)** were also analyzed in the same conditioned media by ELISA. N=3. * P<0.05, ** P<0.01, *** P<0.001.

This shortlist of cytokines was loaded on the STRING database for network analysis, and the resulting network (showing only connections with interaction score >0.7) confirmed the strict relations among them. Moreover, K-means clustering on the functional network evidenced 2 main clusters (Figure 5B). The Gene Ontology analysis on the STRING database returned a long list of significant terms involved. A selection of highly significant GO terms, with high strength (>1) and involving at least 4 counts in the network, is reported in Supplemental Table 3. Representative network images evidencing the molecules involved in a selection of GO terms of interest (concerning features of inflammation, cell migration, ECM synthesis, and vascular diseases) are reported in Supplemental Figure 2.

The differences revealed by the protein array analysis were validated by ELISA, or RT-qPCR for the corresponding mRNAs when the protein concentration was below detection limit. A significantly reduced concentration of secreted adiponectin was confirmed in the supernatant of cells previously treated with both HNBC and TCC serum lots versus NS serum-treated cells (Figure 5C). In addition, significantly reduced levels of *CHI3L1* were confirmed by RT-qPCR in CSCs from both HNBC and TCC treatment groups versus NS control (Figure 5D); significantly increased expression levels of *CRP* were confirmed in TCC serum-treated cells versus NS control, as well (Figure 5E).

Additionally, we quantified the release of H_2_O_2_ and NO in the conditioned media, as indicators of oxidative stress and pro-angiogenic property, respectively. We detected a significantly higher concentration of H_2_O_2_ in conditioned media from cells pre-treated with the TCC serum lot versus the NS lot (Figure 5F), and a significant reduction in NO concentration released by CSCs treated with TCC and HNBC serum lots, both compared to the NS control (Figure 5G). Notably, H_2_O_2_ release was higher from cells previously cultured in the HNBC serum lot compared to the NS control, albeit to a significantly lower concentration compared to CSCs treated with the TCC lot.

### Functional paracrine features after treatment with serum from TCC and HNBC smokers

We then assessed the paracrine effects of CSC-conditioned media concerning support to endothelial cells for angiogenesis, and to cardiomyocyte for cell viability (Figure 6A). The angiogenic assay on Matrigel with HUVECs evidenced a significant reduction in the pro-angiogenic features of conditioned media from cells pre-treated with both TCC and HNBC serum lots, compared to the NS lot (Figure 6B). Several indexes were evaluated, including number of nodes, master segments and meshes, and the total mesh area (Figure 6C). Moreover, conditioned media were used to culture neonatal rat ventricular myocytes (NRVMs), whose viability was monitored by MTS assay. Not surprisingly, the composition of the media for conditioning (0.1% FBS in IMDM) was not optimal for NRVMs, and accelerated progressive cell death of the primary culture, compared to the positive experimental control (Figure 6D). Nonetheless, media conditioned by CSCs previously exposed to the NS serum lot was able to significantly delay cell death in the first 48 hours of culture, compared to media conditioned by cells exposed to both TCC and HNBC serum lots (Figure 6E). This difference remained significant after 72 hours in the TCC group versus the NS control. Overall, this data indicates a detrimental effect of circulating molecules in TCC and HNBC smokers on the protective paracrine functions of CSCs, albeit to a different extent for some features concerning HNBC smokers.

**Figure 6.**
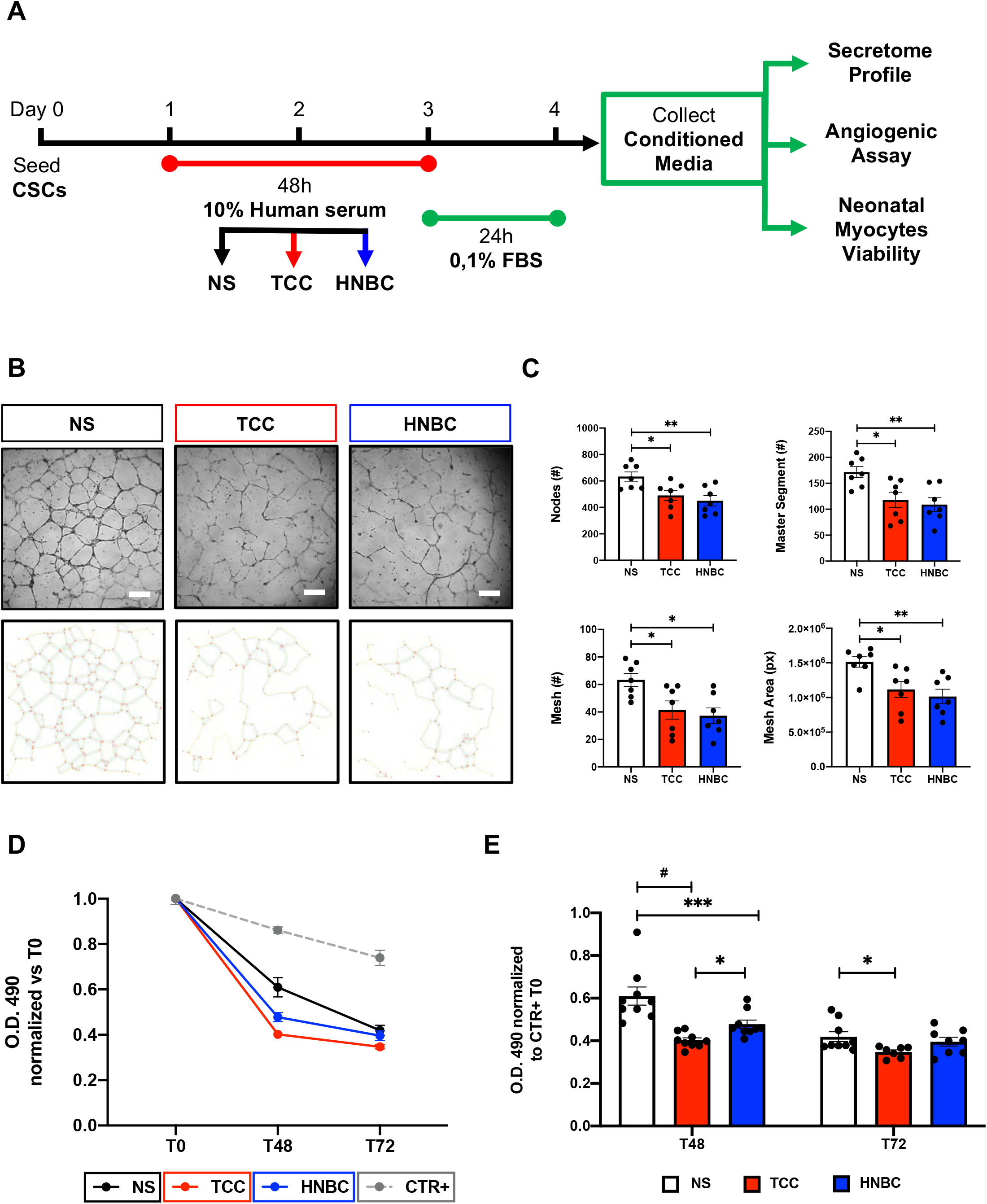
Functional paracrine effects of conditioned media on endothelial cells and cardiomyocytes. **(A)** Experimental design for the functional paracrine experiments with conditioned media. Human umbilical vein endothelial cells (HUVECs) were cultured overnight on Matrigel with conditioned media from the 3 CSC lines previously treated with serum lots from non-smokers (NS), tobacco combustion cigarette (TCC) and heat-not-burn cigarette (HNBC) smokers. Representative images of cultures are shown **(B)**, with quantification of the number of nodes, master segments, meshes, and the mesh area **(C)** in the different conditions. Experiment was performed in triplicate. Two significant outlier replicates were excluded. N=7. Neonatal rat ventricular myocytes (NRVMs) were cultured up to 72 hours with the conditioned media from the 3 CSC lines (each in experimental triplicate), and in EGM2 media as the experimental control (CTR+). Optical Density (OD) values with time of culture are shown **(D)** for the different conditions, together with dot plots bar graph for each single time-point **(E)**. N=9. # = number. * P<0.05, ** P<0.01, *** P<0.001, # P<0.0001. Scale bar = 200 μm.

## DISCUSSION

Tobacco combustion cigarette (TCC) smoking represents a major lifestyle risk factor for cardiovascular diseases, mainly concerning atherosclerosis, which accounts approximately for 25% of cardiovascular deaths (20). The use of novel smoking devices is increasing rapidly worldwide. As of 2018, an estimate of 2.4% among the adult population in the US had used heated tobacco products (source: www.cdc.gov). This proportion appears even higher in the European Union, where heat-not-burn cigarettes (HNBCs) have been introduced on the market earlier, with estimates of 6.5% of adults who had used HNBC products in 2021 (21). These numbers are destined to increase, as more and more people appear to start using HNBCs, alone or in combination with TCCs. Thus, unveiling the exact mechanisms and extent of cardiovascular damage due to chronic HNBC use is of paramount importance to understand and define their prospective impact on health. Many molecular and cellular mechanisms responsible for detrimental cardiovascular effects have been described for TCCs, but there is a significant gap in knowledge about the mechanisms and damage extent related to chronic use of HNBCs.

The patients recruited in the SUR-VAPES Chronic trial as exclusive HNBC chronic smokers represent a one-of-a-kind cohort (10). By using this unique set of smokers’ serum, we have provided data, for the first time, on how CSCs isolated from clinically relevant patients (affected by cardiovascular diseases) can be altered by the circulating molecular fingerprint imposed by chronic HNBC smoke. Our results contribute unprecedented insights on how the myocardial microenvironment may be changed due to chronic HNBC use. Moreover, the present study provides the first basis for a realistic definition of the extent of cardiac insult caused by chronic use of these novel smoking devices. Importantly, our results are obtained on primary atrial resident CSCs, that is a cell population directly involved in chronic atrial fibrosis, which is the main known mechanism involved in the incidence of atrial fibrillation in chronic TCC smokers (2).

Results have shown how the TCC serum lot is less trophic for CSCs to grow as spheroids, compared to serum lots of non-smokers (NS) and HNBC smokers. For this and other phenotypic features, the HNBC serum lot appeared to elicit effects similar to the NS lot, but different from the TCC’s, for example considering the increase in profibrotic CD90+ cells with time in culture, or the stimulation of migration. Concerning the acute effects on gene expression, the serum from HNBC smokers appeared to directly modulate the mRNA levels of markers of fibroblast activation and differentiation (e.g. *COL1A1, THY1, ACTA2*) and of potential contribution to arrhythmias. In fact, the HNBC lot was the only one able to significantly reduce both mRNA and protein levels of *GJ1A/*Cx43 in our conditions, consistently with a pro-fibrotic and pro-arrhythmogenic shift of the stromal population. Conversely, the serum of TCC smokers was the only one capable of inducing the modulation of pro-inflammatory genes (e.g. *IL6, IL8*). In this regard, we can speculate that both TCC and HNBC sera can induce cardiac fibrosis, but the one from TCC smokers appears to induce a more evident pro-inflammatory profile, possibly linked to a reduced trophic support and to even stronger pathogenetic drive.

The difference between TCC and HNBC serum lots was also evident in the profiles of secreted factors by CSCs, where exposure to the serum of TCC smokers was specifically associated to higher release of many immune activating cytokines by CSCs (e.g. CRP, CXCL4, CD40L, C5/C5A, IL17A). Nonetheless, the secreted levels of many protective molecules (e.g. adiponectin, NO, CHI3L1) were similarly reduced in CSC-conditioned media after exposure to both HNBC and TCC serum lots, compared to NS control. Interestingly, the impairment in paracrine support to endothelial cells from CSCs appeared similar after treatment with both HNBC and TCC sera, compared to NS control. This was consistent with NO release levels, although VEGF mRNA and protein levels appeared significantly reduced only with treatment of the HNBC serum lot in our conditions. Conversely, the paracrine pro-survival support to NRVMs appeared more compromised in CSCs exposed to the TCC serum lot, while treatment with the HNBC lot performed in between the TCC and NS groups. These results point to the reduction of beneficial paracrine effects by the cardiac stroma, and are in line with studies reporting a worst prognosis after myocardial infarction in smokers, associated with enhanced acute inflammation and higher incidence of infarct zone hemorrhage (22).

To the best of our knowledge, only one study so far has investigated the effects of human serum derived from modified risk product smokers on the phenotype of cells of cardiovascular interest, focusing on human induced pluripotent stem cell (iPS)-derived endothelial cells (12). Authors of that study had mixed sera of two vaping e-cigarette smokers with those of two dual e-cigarette/TCC smokers, comparing this mixed pool with sera of TCC smokers that were 10 years older, on average. Thus, the investigation on sera from HNBC smokers has never been performed previously. Moreover, the serum lots obtained from the SUR-VAPES Chronic trial used in the present report correspond to a unique homogenous cohort of 60 matched subjects, among which the HNBC group included only exclusive chronic HNBC smokers, thus strengthening the reproducibility and significance of our results. Furthermore, here we studied the response of primary human CSCs freshly isolated from atrial tissue of clinically relevant donors, that is patients with cardiovascular diseases.

### Study limitations

One of the limitations of our study is that HNBC smokers from the SUR-VAPES Chronic trial were all former TCC users, although this bias could be mitigated by the significant wash-out lag of 1,5 years, on average, since the switch to exclusive HNBC use.

## CONCLUSIONS

In conclusion, we provide for the first time data supporting the potential trigger of cardiac fibrosis in chronic HNBC smokers. The results here collected show how human CSCs exposed to the circulating molecules of exclusive HNBC chronic smokers display a shift towards a fibrotic phenotype, with reduced release of beneficial paracrine factors and impaired capacity to support angiogenesis. Interestingly, these effects appear somehow different from those generated by the serum of chronic TCC smokers, inducing intermediate features on the phenotype of CSCs. In fact, the serum of TCC smokers caused a much more evident pro-inflammatory profile in CSCs, with significant reduction of migration capacity, angiogenic support, as well as pro-survival signals to cardiomyocytes. Interestingly, the effects of the HNBC serum on those properties of CSCs were either unchanged, or still affected but to a lower extent. The only distinctive and strong effect in our conditions appeared to be the reduction of connexin 43 expression elicited by exposure to HNBC smokers’ serum, raising a worrying alarm against possible arrhythmogenic effects of chronic HNBC use. Overall, results suggest that the specific molecular profile in the serum of HNBC smokers may act as an important trigger for cardiac fibrosis through the altered functional and paracrine phenotype of resident stromal cells, albeit to a different extent when compared to traditional TCC smokers.

### Perspectives

#### Competency in Medical Knowledge

Tobacco smoking represents a significant risk factor for many cardiovascular diseases, including cardiac fibrosis and atrial fibrillation. The use of novel smoking devices, such as heat-not burn cigarettes, is raising exponentially, and their impact on health is still uncertain.

#### Translational outlook 1

The circulating molecules in the serum of chronic exclusive smokers of heat-not burn cigarettes appear to trigger a pro-fibrotic specification in human atrial stromal cells.

#### Translational outlook 2

The chronic use of HNBCs could increase the risk of cardiac fibrosis development.

## ABBREVIATIONS LIST

CSCs: Cardiac stromal cells
NS: Non-smoker
TCC: Tobacco combustion cigarettes
HNBCs: Heat-not burn cigarettes
MRP: Modified risk product
ECM: Extracellular matrix
NRVMs: neonatal rat ventricular myocytes
iPSCs: induced pluripotent stem cell

## FIGURE LEGENDS

**Supplemental figure 1.**
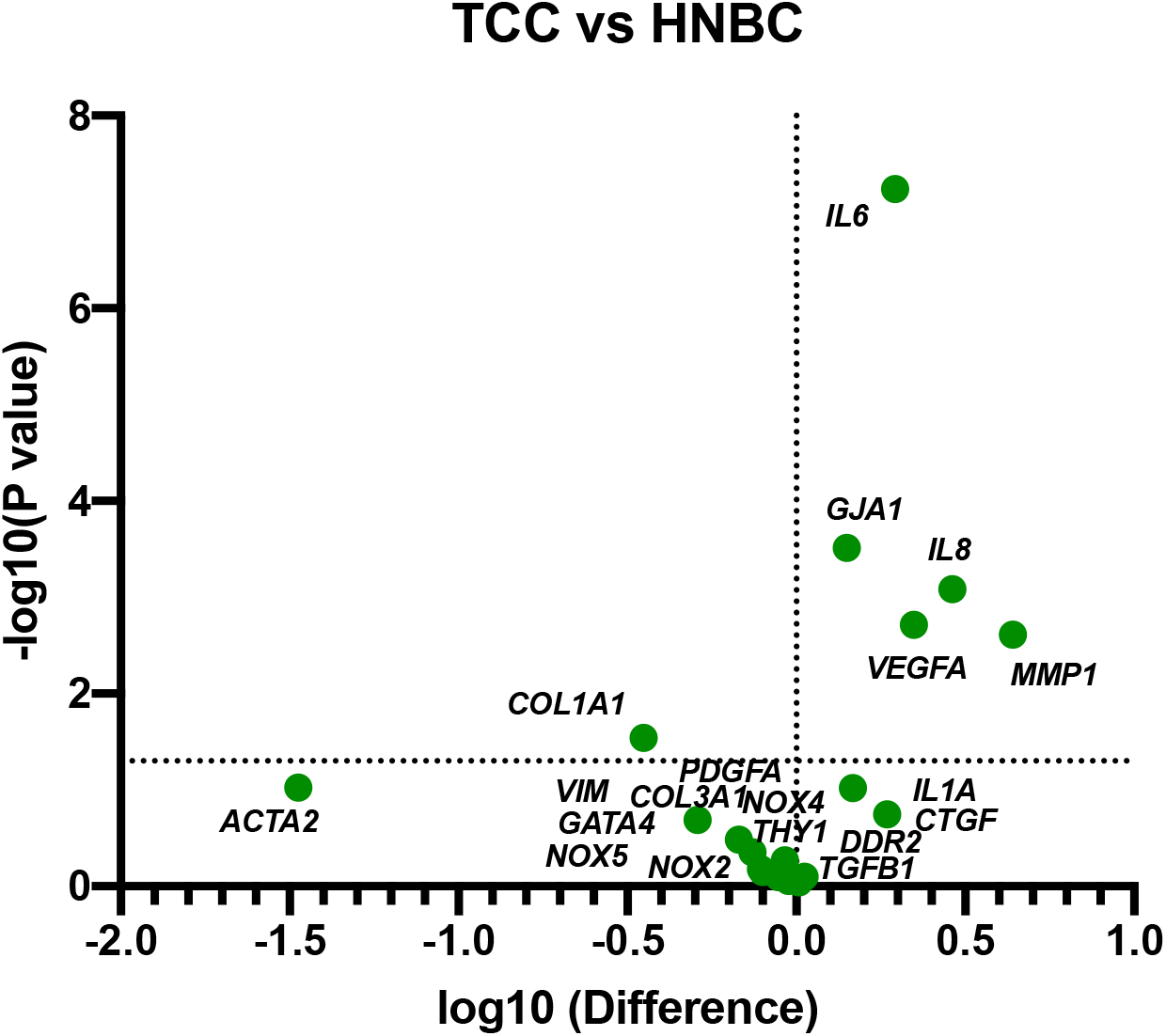

**Supplemental Figure 2.**
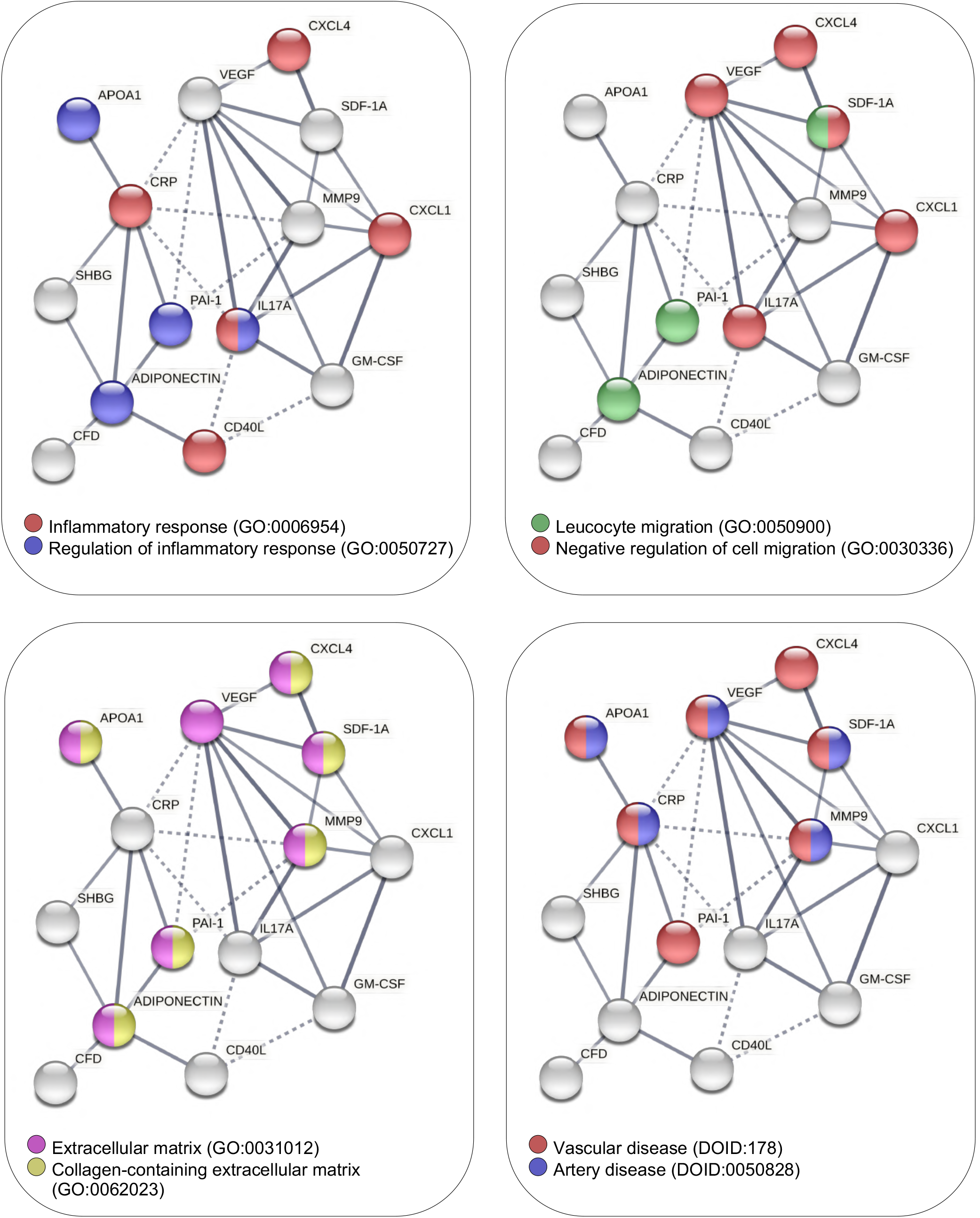

