## Supplemental material for "Investigating the risk of cardiac fibrosis due to heat-not-burn cigarettes through human cardiac stromal cells"

**Content of this appendix:**

| | |
|---|---|
| **Supplemental Methods** | **page 2** |
| **Supplemental Tables** | **page 5** |
| **Supplemental Figures** | **page 12** |

### Supplemental Methods

**Flow cytometry.** CSCs from the first explant harvest and at passage 1 were used for all donors. Cells from semi-confluent cultures were harvested with trypsin-EDTA and stained with CD90-FITC (Dianova, AS02) using 300ng antibody in 100µl staining buffer (PBS with 2% FBS) per sample. All acquisitions were performed using a BD FACS-Aria II cytometer equipped with DIVA software (BD Biosciences), which was also used to calculate the compensation parameters. All flow cytometry data were analyzed with FlowJo software (FlowJo LLC).

**RNA extraction and RT-qPCR.** Total RNA was extracted using the miRNeasy Micro Kit (Qiagen) and quantified using Nanodrop spectrophotometer (Thermo Fisher Scientific). cDNA was synthesized from 0.5µg RNA with the High-Capacity cDNA Reverse Transcription Kit (Thermo Fisher Scientific). Real-time qPCR was performed to assess gene expression using Power SYBR Green PCR Master Mix (Thermo Fisher Scientific) and standard thermocycling conditions, according to the manufacturer's protocol. The relative ratio versus reference gene was calculated using the comparative Ct method (2^-ΔCt). The set of genes analyzed, and the primers sequences are listed in Supplemental Table 4. Hypoxanthine Phosphoribosyltransferase 1 (HPRT1) was selected as the housekeeping gene according to the Bestkeeper spreadsheet macro (freely available at: www.gene-quantification.de). The PCR data was analyzed (2^-ΔCt for the retrospective analysis, or fold change versus NS for the in vitro treatments), and Volcano Plots were generated by multiple t-test analysis using GraphPad Prism 8 software (GraphPad Software).

**Spheroid assay.** For the spheroid forming assay, 1,5x10^4 CSCs per well were plated in 24-well plates coated with Poly-D-Lysine (Corning), in CEM supplemented with 10% human serum from the NS, TCC, or HNBC lots. Whole well images were captured after 7 days of culture with a Nikon Eclipse Ti microscope equipped with NIS-Elements AR 4.30.02 software (Nikon Corporation). Images were analyzed using ImageJ software exploiting the plugins for particle counts and area measurement.

**Immunostaining and Fluorescence Microscopy Analysis.** Cells were fixed for 10 minutes with 4% paraformaldehyde at 4°C, and permeabilized with 0.1% Triton X-100 (Sigma-Aldrich) in PBS with 1% BSA. Nonspecific antibody binding sites were blocked with 10% goat serum (Sigma-Aldrich) in PBS, before overnight incubation at 4°C with primary antibody anti-CX43 (MAB306, Millipore). After thorough washing, slides were incubated for 2 hours at room temperature with

Alexa-conjugated secondary antibody (AF Goat α-mouse 488, A-11001, Thermo-Fisher) and Hoechst nuclear dye (Thermo-Fisher). Slides were mounted in Vectashield (VectorLabs). Image capture was performed on a Nikon Eclipse Ni microscope equipped with VICO system, and average fluorescence intensity per cell area was calculated by semi-automatic methods using NIS-Elements AR 4.30.02 software with a 20X objective (Nikon Corporation).

**Scratch assay**. CSCs (10^5 per well) were plated in 12-well plates coated with Fibronectin (Corning) in CEM 10% FBS. The scratch was performed after 24 hours from plating: cells were washed with PBS and then cultured with CEM supplemented with 10% human serum derived from NS, TCC, or HNBC serum lots. Images were captured after 6, 8 and 10 hours with a Nikon Eclipse Ti fluorescence microscope, equipped with motorized stage and NIS-Elements AR 4.30.02 software (Nikon Corporation). Images were analyzed by ImageJ software (1) using an automatic macro for scratch area measurement.

**Cytokine Array for secretome profiling.** CSCs were pre-treated for 48 hours with CEM supplemented with 10% human serum derived from NS, TCC, or HNBC serum lots. After thorough washing, cells were cultured for the following 24 hours with CEM 0,1% FBS for conditioned media (CM) collection. Media were gently aspirated and centrifuged at 2000rcf for 5 minutes to remove cells and debris, and then stored at -80°C until analysis. Media were assayed using the Proteome Profiler Human XL Cytokine Array (R&D Systems), according to the manufacturer's instructions. Optical density (OD) of membranes was quantified by the ChemiDocXRS+ Imager (Bio-Rad), and densitometric analysis was performed using Image Lab software (Bio-Rad). The data obtained (log2-trasformed densitometric quantification of each dot) was plotted as a heatmap generated using the pheatmap R package (GNU Project). Functional association network was created using the STRING database (ELIXIR Core Data Resources) selecting the "experiments" and "database" as interaction sources, and high confidence (0.70) as the minimum interaction score.

**ELISA assays.** Adiponectin concentration was evaluated using the specific ELISA kit (human adiponectin Elisa kit, Boster Immunoleader), following the manufacturer's protocol. Absorbance at 450nm was recorded immediately after the end of all steps using the Varioskan Lux multimode microplate reader (Thermo Fisher Scientific). The data were analyzed using the Skanlt software (Thermo Fisher Scientific). $H_2O_2$ and NO release were also quantified by ELISA, as previously described and according to the manufacturers' instructions (2), by a Colorimetric Detection Kit (Arbor Assays) and expressed as μmol/L.

**Angiogenesis assay.** The tube forming assay was performed as previously described (2,3), with human umbilical vein endothelial cells (HUVECs) routinely cultured with standard protocols in EGM2 media (Lonza). Cells were plated 1,5x10^4 per well for 16 hours on matrigel-coated 96-well plates (Growth Factor Reduced Matrigel Matrix Phenol Red Free, BD) in presence of the CSC-CMs collected in CEM 0.1% FBS. An automated scan of each well was acquired with a 4X objective by Nikon Eclipse TI inverted microscope with motorized stage (Nikon Corporation). Quantification of the number of nodes, number of master segments, number of meshes, and mesh area were performed by ImageJ software (1) and the Angiogenesis plugin (NIH).

**MTS Cell viability assay.** MTS cell viability assay was performed to evaluate NRVMs viability using Cell Titer 96® Aqueous Non-Radioactive Cell Proliferation Assay (MTS) (Promega). NRVMs were isolated from 2 day-old Sprague Dawley rat pups by the Neonatal Cardiomyocyte Isolation Kit, Rat, on a gentleMACS Dissociator (Miltenyi Biotec), according to the manufacturer's protocols. NRVMs were plated in 96-well plates (1x10^3 per well) in 100 µl Cardio Medium DMEM/F-12 (Sigma, D0547) supplemented with 0.72 g/L glucose (Sigma, G5400), 0,33 g/L sodium pyruvate (Thermo Fisher Scientific, BP356), 0.017 g/L ascorbic acid (Gibco, 13080-23), 2 µl selenite 0.2 M (Sigma, S5261), 0.004 g/L transferrin (Sigma, T3309), 2 g/l BSA fraction V (VWR, 0332), 3.57 g/l HEPES (VWR, 0511), 2.43 g/l sodium bicarbonate (Sigma, S6014) and 10 ml penicillin-streptomycin (Gibco, 15070063). After 24 hours, NRVMs were incubated with conditioned media, or Cardio Medium as the positive experimental control. NRVMs viability was measured 48 and 72 hours after treatment adding 20 µl of combined MTS/PMS Solution to each well containing cells in 100 µl of medium, and incubating for 2 hours. The absorbance at 490 nm was recorded using Varioskan™ LUX Multimode Reader (Thermo Fisher Scientific). The data were analyzed using the Skanlt software (Thermo Fisher Scientific).

**Supplemental Table 1. Clinical characteristics of the study population of biopsy donors for the retrospective analysis.** Comparative analysis of anthropometric and clinical characteristics of smokers and non-smokers patients, from which CSCs had been isolated and characterized.

| Feature* | All patients, N=38 | Smokers, N=18 | Non-smokers, N=20 | Significance# |
|---|---|---|---|---|
| **Age, years** | 68 (54; 79) | 64 (54; 78) | 65 (63; 80) | 0.01 |
| **Gender, male (%)** | 25 (65%) | 16 (88%) | 9 (45%) | 0.006 |
| **BMI, kg/m$^2$** | 26.5 (19; 33) | 25.4 (19;29.7) | 26.5 (20.8; 34) | 0.09 |
| **BSA, m$^2$** | 1.7 (1.4; 2.2) | 1.9 (1.6; 2.1) | 1.6 (1.4; 2.2) | 0.02 |
| **Systolic BP, mmHg** | 117 (90; 190) | 105 (110; 150) | 122 (90; 190) | 0.72 |
| **Diastolic BP, mmHg** | 134 (60; 120) | 61 (60; 80) | 75 (60; 120) | 0.32 |
| **Heart rate, bpm** | 67 (55; 100) | 61 (50; 100) | 68 (50; 98) | 0.74 |
| **Atrial fibrillation, N (%)** | 7 (18.4%) | 1 (5.5%) | 4 (20%) | 0.34 |
| **Previous myocardial infarction, N (%)** | 5 (23%) | 5 (28%) | 4 (20%) | 0.71 |
| **Recent (<30 days) myocardial infarction, N (%)** | 11 (29%) | 7 (39%) | 4 (20%) | 0.29 |
| **Diabetes, N (%)** | 18 (47.3%) | 6 (33.3%) | 6 (30%) | 0.99 |
| **Metabolic Syndrome, N (%)** | 12 (31.5%) | 5 (27.7%) | 7 (35%) | 0.73 |

*Reported as median (1st; 3rd quartile) for continuous variables and count/total (percentage) for categorical variables. #Computed with unpaired Mann-Whitney U test for continuous variables and Fisher exact test for categorical variables.

BMI: body mass index. BSA: body surface area. BP: blood pressure. BPM: beats per minute.

**Supplemental Table 2. Surgical characteristics of the study population of biopsy donors for the retrospective analysis.** Comparative analysis of surgical characteristics of smokers and non-smokers patients, from which CSCs had been isolated and characterized.

| Feature* | All patients, N=38 | Smokers, N=18 | Non-smokers, N=20 | Significance# |
|---|---|---|---|---|
| **STS mortality risk score** | 1.9 (0.8; 2.6) | 2.1 (0.2; 8.7) | 1.8 (7.2; 0.4) | 0.69 |
| **Euroscore II** | 3.4 (0.5; 12) | 2.2 (0.5; 4.9) | 4.5 (11.8; 1.01) | 0.03 |
| **Coronary by-pass surgery, N (%)** | 23 (60.5%) | 17 (94.4%) | 11 (55%) | 0.009 |
| **Heart valve surgery, N (%)** | 9 (23.6%) | 5 (27.7%) | 10 (50%) | 0.19 |
| **Coronary by-pass + heart valve surgery, N (%)** | 6 (15.7%) | 3 (16.6%) | 3 (15%) | 0.99 |
| **Other surgery, N (%)** | 1 (2.6%) | 0 (0%) | 1 (5%) | 0.99 |
| **CPB time, min** | 140 (74; 232) | 138 (74; 220) | 148 (89; 232) | 0.15 |
| **Cross clamp time, min** | 107 (46; 207) | 107 (46; 182) | 112 (53; 207) | 0.36 |

*Reported as median (1st; 3rd quartile) for continuous variables and count/total (percentage) for categorical variables. #Computed with unpaired Mann-Whitney U test for continuous variables and Fisher exact test for categorical variables.

STS: society of thoracic surgery. CBP: cardiopulmonary bypass.

**Supplemental table 3. Gene ontology terms associated with the cytokine STRING network.**

| **Biological Process** | | | | | |
|---|---|---|---|---|---|
| **term ID** | **term description** | **observed gene count** | **strength** | **FDR** | **matching proteins network** |
| **GO:0010743** | Regulation of macrophage derived foam cell differentiation | 4 | 2.08 | 2.38e-05 | crp, cxcl4, gm-csf, adiponectin |
| **GO:0097530** | Granulocyte migration | 4 | 1.6 | 0.00050 | cxcl4, il17a, cxcl1, vegf |
| **GO:0002688** | Regulation of leukocyte chemotaxis | 5 | 1.57 | 7.44e-05 | mif, pai-1, c5, sdf-1a, vegf |
| **GO:0002576** | Platelet degranulation | 5 | 1.56 | 8.76e-05 | pai-1, apoa1, cxcl4, cfd, vegf |
| **GO:0030595** | Leukocyte chemotaxis | 4 | 1.42 | 0.0015 | cxcl4, cxcl1, sdf-1a, vegf |
| **GO:0030856** | Regulation of epithelial cell differentiation | 4 | 1.39 | 0.0019 | pai-1, mmp9, adiponectin, vegf |
| **GO:0032680** | Regulation of tumor necrosis factor production | 4 | 1.39 | 0.0019 | mif, cxcl4, il17a, adiponectin |
| **GO:1905952** | Regulation of lipid localization | 4 | 1.39 | 0.0019 | mif, apoa1, crp, adiponectin |
| **GO:2001234** | Negative regulation of apoptotic signaling pathway | 6 | 1.38 | 5.17e-05 | mif, pai-1, cxcl4, gm-csf, mmp9, sdf-1a |
| **GO:0060326** | Cell chemotaxis | 5 | 1.36 | 0.00046 | c5, cxcl4, cxcl1, sdf-1a, vegf |
| **GO:0006959** | Humoral immune response | 6 | 1.31 | 0.00012 | c5, crp, cxcl4, cfd, cxcl1, sdf-1a |
| **GO:0022408** | Negative regulation of cell-cell adhesion | 4 | 1.31 | 0.0033 | apoa1, sdf-1a, adiponectin, vegf |

| | | | | | |
|---|---|---|---|---|---|
| **GO:0071222** | Cellular response to lipopolysaccharide | 4 | 1.3 | 0.0034 | pai-1, cxcl4, gm-csf, cxcl1 |
| **GO:0050731** | Positive regulation of peptidyl-tyrosine phosphorylation | 4 | 1.28 | 0.0041 | mif, gm-csf, adiponectin, vegf |
| **GO:0050900** | Leukocyte migration | 6 | 1.25 | 0.00022 | mif, cxcl4, il17a, cxcl1, sdf-1a, vegf |
| **GO:0030336** | Negative regulation of cell migration | 5 | 1.25 | 0.00092 | mif, pai-1, c5, sdf-1a, adiponectin |
| **GO:0043406** | Positive regulation of map kinase activity | 5 | 1.25 | 0.00096 | mif, c5, gdf-15, cd40l, vegf |
| **GO:0007162** | Negative regulation of cell adhesion | 5 | 1.22 | 0.0011 | pai-1, apoa1, sdf-1a, adiponectin, vegf |
| **GO:0006954** | Inflammatory response | 9 | 1.21 | 1.62e-06 | mif, c5, crp, chi3l1, ptx3, cxcl4, il17a, cd40l, cxcl1 |
| **GO:0001819** | Positive regulation of cytokine production | 8 | 1.21 | 9.40e-06 | mif, pai-1, c5, cxcl4, gm-csf, il17a, cd40l, adiponectin |
| **GO:0043405** | Regulation of map kinase activity | 6 | 1.21 | 0.00029 | mif, c5, gdf-15, cd40l, adiponectin, vegf |
| **GO:0071902** | Positive regulation of protein serine/threonine kinase activity | 6 | 1.21 | 0.00031 | mif, c5, gdf-15, cd40l, adiponectin, vegf |
| **GO:0010951** | Negative regulation of endopeptidase activity | 4 | 1.18 | 0.0084 | pai-1, c5, mmp9, vegf |
| **GO:0045637** | Regulation of myeloid cell differentiation | 4 | 1.16 | 0.0097 | cxcl4, gm-csf, il17a, adiponectin |
| **Cellular Component** | | | | | |
| **GO:0031093** | Platelet alpha granule lumen | 4 | 1.74 | 0.00015 | pai-1, cxcl4, cfd, vegf |

| | | | | | |
|---|---|---|---|---|---|
| **GO:0034774** | Secretory granule lumen | 9 | 1.41 | 1.48e-08 | mif, pai-1, apoa1, chi3l1, ptx3, cxcl4, cfd, cxcl1, vegf |
| **GO:0031012** | Extracellular matrix | 9 | 1.2 | 5.00e-07 | pai-1, apoa1, chi3l1, ptx3, cxcl4, mmp9, sdf-1a, adiponectin, vegf |
| **GO:0062023** | Collagen-containing extracellular matrix | 6 | 1.15 | 0.00039 | pai-1, apoa1, cxcl4, mmp9, sdf-1a, adiponectin |
| **GO:0030141** | Secretory granule | 10 | 1.04 | 1.32e-06 | mif, pai-1, apoa1, chi3l1, ptx3, cxcl4, cfd, mmp9, cxcl1, vegf |
| **Disease** | | | | | |
| **DOID:178** | Vascular disease | 7 | 1.47 | 1.24e-05 | pai-1, apoa1, crp, cxcl4, mmp9, sdf-1a, vegf |
| **DOID:0050828** | Artery disease | 5 | 1.6 | 0.00037 | apoa1, crp, mmp9, sdf-1a, vegf |
| **DOID:612** | Primary immunodeficiency disease | 6 | 1.12 | 0.0031 | mif, c5, crp, il17a, cd40l, sdf-1a |
| **DOID:848** | Arthritis | 4 | 1.54 | 0.0031 | mif, c5, crp, il17a |

**Supplemental table 4. Sequence of all Human primers used for real-time PCR.**

| **Human primer name** | **Sequence** | **Human primer name** | **Sequence** |
|---|---|---|---|
| **CRP Forward** | TCGACCCGTGGGTACAGTAT | **PDGFa Forward** | TTTGGACACCAGCCTGAGAG |
| **CRP Reverse** | ATTTCAGACCCACCCACTGT | **PDGFa Reverse** | AGACAGCGGGGACAGCTT |
| **CHI3L1 Forward** | TCTACGGCATGCTCAACACA | **MMP1 Forward** | ACAACTGCCAAATGGGCTTG |
| **CHI3L1 Reverse** | ACTTGATGAAAGTCCGGCGA | **MMP1 Reverse** | TGTCCTTGGGGTATCCGTGT |
| **NOX2 Forward** | CTCACTGGCTGGGATGAGTC | **CTGF Forward** | AGGAGTGGGTGTGTGACGA |
| **NOX2 Reverse** | CATTATCCCAGTTGGGCCGT | **CTGF Reverse** | CCAGGCAGTTGGCTCTAATC |
| **NOX4 Forward** | AACCAAGGGCCAGAGTATCA | **DDR2 Forward** | AGAGGCCAGCTGTTTTTGAG |
| **NOX4 Reverse** | GGATAAGGCTGCAGTTGAGG | **DDR2 Reverse** | AGATAGGCAGCAGCAGGAAC |
| **NOX5 Forward** | CATCGATGTGTGTGCACGGC | **GATA4 Forward** | GTTTTTTCCCCTTTGATTTTTGATC |
| **NOX5 Reverse** | ATCCGGGTCAATGGAGCCAC | **GATA4 Reverse** | AACGACGGCAACAACGATAAT |
| **COL1A1 Forward** | CACACGTCTCGGTCATGGTA | **VEGF Forward** | AGCTGCGCTGATAGACAT |

| | | | |
|---|---|---|---|
| **COL1A1 Reverse** | AAGAGGAAGGCCAAGTCGAG | **VEGF Reverse** | CTACCTCCACCATGCCAA |
| **COL3A1 Forward** | CATGCCCTACTGGTCCTCAG | **VIMENTIN Forward** | ACCCACTCAAAAAGGACACTTC |
| **COL3A1 Reverse** | ATAGCCTGCGAGTCCTCCTA | **VIMENTIN Reverse** | GGTCATCGTGATGCTGAGAA |
| **IL6 Forward** | GGTACATCCTCGACGGCATCT | **THY1 Forward** | CAGCGGAAGACCCCAGT |
| **IL6 Reverse** | GTGCCTCTTTGCTGCTTTCAC | **THY1 Reverse** | CGTTAGGCTGGTCACCTTCT |
| **IL8 Forward** | CTTGGCAGCCTTCCTGATTT | **TGF-b Forward** | CGACTCGCCAGAGTGGTTAT |
| **IL8 Reverse** | TTCTTTAGCACTCCTTGGCAAAA | **TGF-b Reverse** | CTAAGGCGAAAGCCCTCAAT |
| **aSMA Forward** | ATGAAGATCCTGACTGAGCG | **HPRT1 Forward** | TCCTCCTCCTGAGCAGTCA |
| **aSMA Reverse** | GCAGTGGCCATCTCATTTTC | **HPRT1 Reverse** | ACCCTTTCCAAATCCTCAGC |

**Supplemental Figure 1**

**Gene expression analysis of CSCs after treatment with serum lots of smokers**.
The relative gene expression levels of several markers, with corresponding significance values, is plotted as volcano plot for the comparison of tobacco combustion cigarette (TCC)- versus heat-not burn cigarette (HNBC)-serum lots treatment of CSCs. Difference in expression is plotted as log10 fold-change of 2^(-ΔCt).

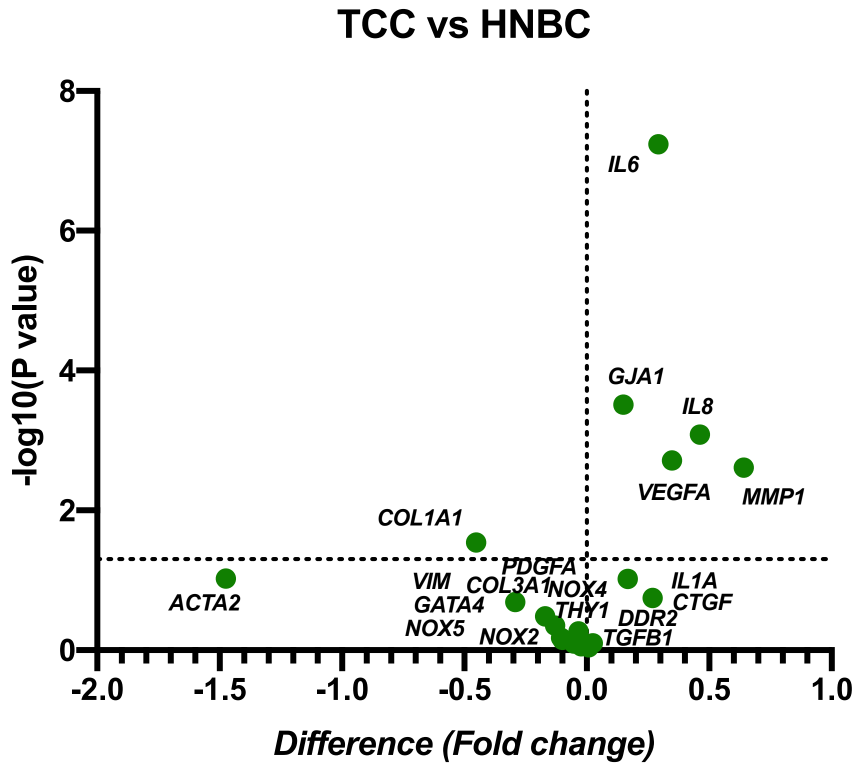

**Supplemental figure 2**

**Gene ontology terms associated with the cytokine STRING network**. The network created by STRING database is colored to highlight cytokines belonging to a selection of Gene Ontology terms of interest, among those with strength >1, and encompassing at least 4 cytokines of the shortlist.

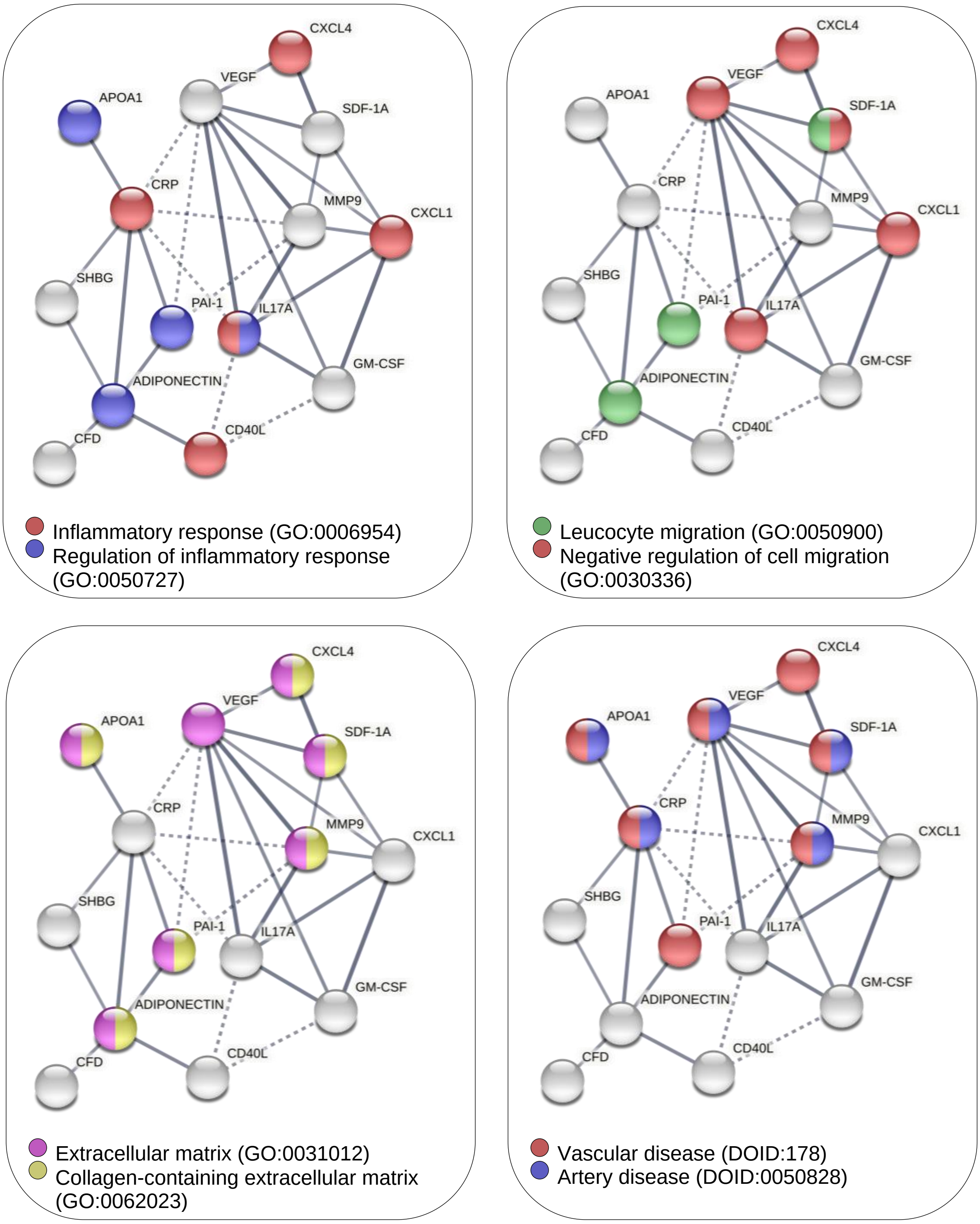
